# The bacterial microbiome of the coral skeleton algal symbiont *Ostreobium* shows preferential associations and signatures of phylosymbiosis

**DOI:** 10.1101/2022.12.13.520198

**Authors:** B.L.D. Uthpala Pushpakumara, Kshitij Tandon, Anusuya Willis, Heroen Verbruggen

**Author notes:** Corresponding Author: email –.

## Abstract

*Ostreobium*, the major algal symbiont of the coral skeleton, remains understudied despite extensive research on the coral holobiont. The enclosed nature of the coral skeleton might reduce the dispersal and exposure of residing bacteria to the outside environment, allowing stronger associations with the algae. Here, we describe the bacterial communities associated with cultured strains of 5 *Ostreobium* clades using 16S rRNA sequencing. We shed light on their likely physical associations by comparative analysis of three datasets generated to capture (1) all algae associated bacteria (2) enriched tightly attached and potential intracellular bacteria and (3) bacteria in spent media. Our data showed that while some bacteria may be loosely attached, some tend to be tightly attached or potentially intracellular. Although colonised with diverse bacteria, *Ostreobium* preferentially associated with 34 bacterial taxa revealing a core microbiome. These bacteria include taxa known as nitrogen cyclers, polysaccharide degraders, sulphate reducers, antimicrobial compound producers, methylotrophs and vitamin B12 producers. By analysing co-occurrence networks of 16S rRNA datasets from *Porites lutea* and *Paragoniastrea australensis* skeleton samples, we show that the *Ostreobium*-bacterial associations present in the cultures are likely to also occur in their natural environment. Finally, our data show significant congruence between the *Ostreobium* phylogeny and the community composition of its tightly associated microbiome, largely due to the phylosymbiotic signal originating from the core bacterial taxa. This study offers insight into the *Ostreobium* microbiome and reveals preferential associations that warrant further testing from functional and evolutionary perspectives.

## Introduction

Bacterial interactions with algae are important due to their influence on ecosystem productivity. Through culture-based experiments [1], [2], omics approaches [3], [4], and microbial network studies [5], [6], a range of algal associations with bacteria have been uncovered. The algal phycosphere, an area around the algal cell rich in photosynthetic exudates, is home to diverse bacteria [7], [8] with host interactions that range from mutualistic to parasitic [9], [10]. Organic carbon and inorganic nutrient exchanges are the most common interactions observed between algae and bacteria [11], [12], while more specialised interactions include provision of B vitamins [13], [14], iron chelating siderophores [15], growth promoting hormones [16] by bacteria.

Tightly associated algal-bacterial systems provide an opportunity to investigate the functional nature and evolutionary basis of algal-bacterial interactions. Phylosymbiosis captures the correlation between the host phylogeny and the relationships of microbial communities associated with those hosts [17]. Both deterministic processes, such as influence on community composition by host traits, and random processes, like changes in microbial community dispersal and host geographical ranges, may lead to phylosymbiosis [18], [19]. It has been observed in a number of terrestrial systems [20]–[22] and is now gaining attention in marine hosts due to major studies on scleractinian corals [23] and sponges [24] detecting this pattern.

*Ostreobium*, a siphonous green alga, is an endolithic alga living in marine limestone, and is the principal skeletal alga of corals [25]. It lives in near darkness, variable concentrations of O_2_ and fluctuating pH [26]–[28]. During bleaching events, *Ostreobium* is shown to provide photosynthates to the coral organism helping to keep it temporarily alive [29], [30]. As an endolithic alga, *Ostreobium* lives in a confined environment with its associated bacteria, which likely reduces the frequency of opportunities for bacterial recruitment or exchange. Such an environment may have resulted in evolutionary pressure to conserve associations and co-disperse. In other confined spaces, for example such as animal gut, stronger eco-evolutionary patterns have been observed [31].

Extensive work on different biological components associated with corals are carried out to shed light on the coral holobiont. A large gap remains in our knowledge of microbial associations in the coral skeleton, including with the major skeletal algal symbiont, *Ostreobium*. The aim of this study was to describe the bacterial communities associated with cultured strains from different *Ostreobium* lineages. Using different sample processing protocols, we focus on investigating likely physical associations of bacteria with its algal host. We investigate if *Ostreobium*-bacteria associations present in the cultures are likely to also occur in their natural environment through co-occurrence network analysis. We test whether evolutionary relationships between *Ostreobium* lineages are associated with differences in bacterial community composition, with the prediction that microbial dendrograms built on beta diversity differences would be more congruent with *Ostreobium* phylogeny than expected at random.

## Methods

### Study design, sample preparation and DNA extraction

Cultures of five strains (VRM605, VRM642, VRM644, VRM646, VRM647) representing five clades (P3P14, C, P1K, P4, B3, respectively) of *Ostreobium* were used. Isolation of these strains from skeleton fragments of Great Barrier Reef corals and molecular identification were described previously [32]. To investigate likely physical associations, three sample processing protocols were employed which we will call ‘whole’, ‘media’’ and ‘tight’. All sample processing was conducted on the same day. For the first protocol, *Ostreobium* filaments were collected, placed in sterile centrifuge tubes and snap frozen, thereby characterising all the culture associated bacteria (attached, intracellular and unattached in the culturing media) and the data stemming will be referred to as the ‘whole’ dataset. For the second protocol, to generate ‘media’ dataset, fifty millilitres of spent media from each culture was collected, filtered using 0.22µm and filters were placed in sterile centrifuge tubes and snap frozen. This data was used to identify unattached and contaminating bacteria. The third protocol was aimed at characterising potentially intracellular as well as tightly attached bacteria. Algal filaments were washed serially, three times, in sterile modified f/2 media by vortexing for 15-20 seconds. The washed algal material was then placed in sterile petri dishes with DNA extraction buffer and the filaments were cut with sterile dissecting scissors allowing the cytoplasmic material to flow out. Finally, the green cytoplasmic content was collected, avoiding the cell wall material as much as possible to enhance the potentially intracellular taxa and snap frozen until processing. The data generated from this protocol will be referred to as the ‘tight’ dataset. All the snap frozen samples were stored at -20°C until processing. To test for contaminant bacteria in the algal growth room, sterile culture media (modified f/2) in a culture flask was maintained with the algal cultures. Negative controls for filter units and media used to wash the algal material were also tested. DNA extractions were performed following [33] with modifications. Blank DNA extractions were conducted as negative controls.

### 16S rRNA gene PCR amplification, library preparation, and sequencing

Hyper variable regions of the 16S rRNA-gene, V5-V6, were amplified using the primer pairs: 784F [5’-<u>TCGTCGGCAGCGTCAGATGTGTATAAGAGACAG</u>AGGATTAGATACCCTGGTA -3’], and 1061R [5’-<u>GTCTCGTGGGCTCGGAGATGTGTATAAGAGACAG</u>CRRCACGAG CTGACGAC-3’] [34] using a 2 step PCR protocol. Illumina adapters that were attached to the primers are shown as underlined. In the first PCR round, each reaction contained 10uL of KAPA HiFi HotStart ReadyMix, 0.5uL of each primer (10uM) and 10uL of DNA template. Previously described PCR conditions [35]were used for 1^st^ and 2^nd^ PCR steps. First-PCR conditions were as follows: 95 °C for 3 min; 25 cycles of 98 °C for 20 s, 60 °C for 15 s, 72 °C for 30 s, a final extension at 72 °C for 1 min. In the second PCR round, each reaction contained 10uL GoTaq Green mix, 0.5uL of each custom-made Illumina index (10uM) and 10uL DNA template. PCR conditions were as follows: 95 °C for 3 min; 24 cycles each at 95 °C for 15 s, 60 °C for 30 s, 72 °C for 30 s, a final extension at 72 °C for 7 min. Triplicate PCRs were conducted for each sample and controls. Three PCRs without template DNA were also conducted. A library pool was prepared taking 5uL from each well per plate, cleaned up using beads (80uL beads to 100uL library pool), quality-checked on a TapeStation (model 4200) and sequenced using the Illumina MiSeq platform (2*300 bp paired end reads) at the Walter and Eliza Hall Institute of Medical Research. In total, we sequenced 6 replicates per strain (2 biological replicates * 3 technical replicates) in each protocol.

### 16S rRNA gene analysis in Qiime2

Raw, demultiplexed sequence reads were analysed in QIIME2 v2021.2. Cutadapt was used to remove contaminating primers [36]. Sequence denoising and chimera checking was performed in DADA2 [37] to correct sequencing errors, low quality bases (Q-score < 20), dereplicate and obtain Amplicon Sequence Variants (ASVs). Taxonomy was assigned in QIIME2 against the SILVA database (v132) trained with a naïve Bayes classifier [38], [39]. ASVs identified as mitochondria, chloroplasts and Archaea were filtered. A phylogenetic tree was constructed using the alignment [40] and phylogeny [41] packages. The filtered table with ASV counts, phylogenetic tree, taxonomy classifications table and metadata file were used to perform downstream statistical analyses in RStudio.

### Statistical community analysis

Statistical analyses were performed and graphs produced using R (v4.0.4) [42] using packages phyloseq (v1.32.0) [43], decontam (v1.11.0) [44], microbiome (v1.10.0) [45], vegan (v2.5.7) [46], indicspecies (v1.7.9) [47] and ggplot2 (v3.3.5) [48]. Statistical tests were considered significant at α= 0.05 unless otherwise stated. Contaminant bacteria were identified using R package decontam using default parameters and removed from the dataset. Evenly sampled ASV tables created by rarefying to the lowest read number for a given sample were used to calculate metrics of alpha diversity (observed ASVs, Simpson index, Shannon index) and beta diversity (Bray Curtis, Unifrac, Weighted-Unifrac). Multivariate homogeneity of group dispersions (PERMDISP) was used to check for the effect of group dispersions on PERMANOVA results. The PERMDISP test determines whether two or more groups are homogeneously dispersed in relation to their species in the samples. An insignificant result indicates homogeneous variations and confirms that PERMANOVA results are reliable. Community structure differences (β-diversity) were statistically compared between ‘media’ and ‘whole’, and ‘media’ and ‘tight’ using permutational multivariate analysis of variance (PERMANOVA). Multivariate homogeneity of group dispersions (PERMDISP) was used to check for the effect of group dispersions on PERMANOVA results. Community structure differences between the datasets were visualised with Principal Coordinate Analysis (PCoA). Similarly, alpha diversity data were also analysed to compare the datasets as above by using the non-parametric Kruskal–Wallis test.

An indicator taxa analysis was carried out using indicspecies package to identify bacterial families significantly associated with ‘whole’, ‘tight’ and ‘media’ datasets with 999 permutations to identify taxa that likely represent the loosely attached, tightly attached/intracellular and unassociated/contaminating taxa. The ‘core’ microbiome was calculated to find taxa present among all the strains/clades (abundance =0.001, prevalence =100/100) in the ‘tight’ dataset. The ‘core’ microbiome was calculated at the ASV level and at different taxonomic levels such as phylum and family by aggregating ASVs to the taxonomic level in question. Overall differences in alpha (Kruskal–Wallis test) and beta diversities (PERMANOVA) among the strains were statistically tested for the ‘tight’ dataset as described above.

To analyse phylosymbiosis, an averaged ASV table was generated from the rarefied ASV table by averaging the abundances as described elsewhere [49]. From this table, Bray–Curtis, UniFrac and weighted UniFrac distances were calculated and these beta-diversity matrices were clustered using the UPGMA method (function: hclust(), method= ‘average’). Resulting dendrograms were exported in newick format. To generate the host phylogenetic tree, previously published *tuf*A sequences were used [32]. Sequences were aligned using MUSCLE [50] in Geneious Prime v2020.1.2 and a maximum likelihood host phylogeny built with the IQ-TREE web server with 1000 bootstraps and best-model selection enabled [51]. Host phylogeny and microbial dendrogram were then compared using matching cluster metrics with 10,000 random trees using a previously published Python script [20]. Co-phylogeny plot was created using phytools (v.1.2-0) [52] to compare host and microbial trees.

### Co-occurrence network analysis on *Porites lutea* and *Paragoniastrea australensis* skeletal samples

We constructed co-occurrence networks from *P. lutea* and *P. australensis* skeletal samples using a previously published 16S rRNA gene dataset [28] (SRA accession PRJNA753944) to investigate *Ostreobium*-bacteria associations in natural settings and to determine if those associations can be seen in cultured *Ostreobium*. Bioinformatic analysis was carried out in Qiime2 to obtain ASV tables representing bacterial and chloroplast sequences for each coral species. Any ASV that was present in less than 20% of samples were filtered to remove low prevalent organisms from the correlation analysis. Bacterial taxonomy was assigned using the SILVA database (v132). Chloroplast sequences were reclassified using PhytoRef [53] to identify microalgae in the skeletal samples. Correlation analysis was performed using a previously published shell script [6] in FastSpar. Two individual undirected weighted networks were created using statistically (p<0.05) significant correlations (>0.5) for each coral species by using the igraph [54] package. Networks were visualised in Cytoscape (version 3.8)[55] and clustered using clusterMaker [56] with the MCL clustering algorithm to identify microbial communities. The Network Analyzer plugin [57] was used to compute global network properties of each network.

## Results

### Sequencing overview and bacterial microbiome diversity

To determine likely physical associations, three datasets, ‘whole’, ‘tight’ and ‘media’ were used in this study. We wished to distinguish host associated microbiomes, using ‘whole’ and ‘tight’ datasets, from loosely and unassociated microbiomes determined using the ‘media’ dataset. Sequencing generated 5,526,746 reads across all the algal samples and controls in the “whole’ dataset. After removal of contaminants, 446 bacterial ASVs were observed across the 5 *Ostreobium* strains (Supplementary data file). In ‘tight’ data, sequencing produced 6,619,904 reads across all the algal samples and the controls. After removal of contaminants, 853 ASVs were observed across the strains (Supplementary data file). While in the ‘media’ dataset 5,352,716 reads were generated and had a final ASV count of 416 (Supplementary data file).

Statistically significant community structure differences were evident based on Bray-Curtis distances between ‘media’ and ‘whole’ (Figure 1: a) (PERMANOVA _mediaVSwhole_ = p=0.001, F. Model=7.4907, R^2^= 0.11438) as well as ‘media’ and ‘tight’ datasets (PERMANOVA _mediaVtight_ p=0.001, F. Model=4.9371, R^2^=0.07971) (Figure1: b). Substantially higher ASV richness, evenness and overall Shannon diversity were observed in ‘whole’ and ‘tight’ than ‘media’ data (Figure 1: c). Overall, ‘tight’ data showcased the highest overall alpha diversity. All three alpha diversity metrics were significantly different between ‘media’ and ‘whole’ data (Kruskal-Wallis test: chi-squared _Observed (1)_ =15.774, p = 7.139e-05, chi-squared _Simpson(1)_ =21.414, p = 3.701e-06, chi-squared _Shannon(1)_ =24.093, p = 9.181e-07) and ‘media’ and ‘tight’ data (Kruskal-Wallis test: chi-squared _Observed (1)_ = 32.178, p =1.407e-08, chi-squared _Simpson(1)_ =39.211, p =3.803e-10, chi-squared _Shannon(1)_ =38.455, p = 5.602e-10).

**Figure 1.**
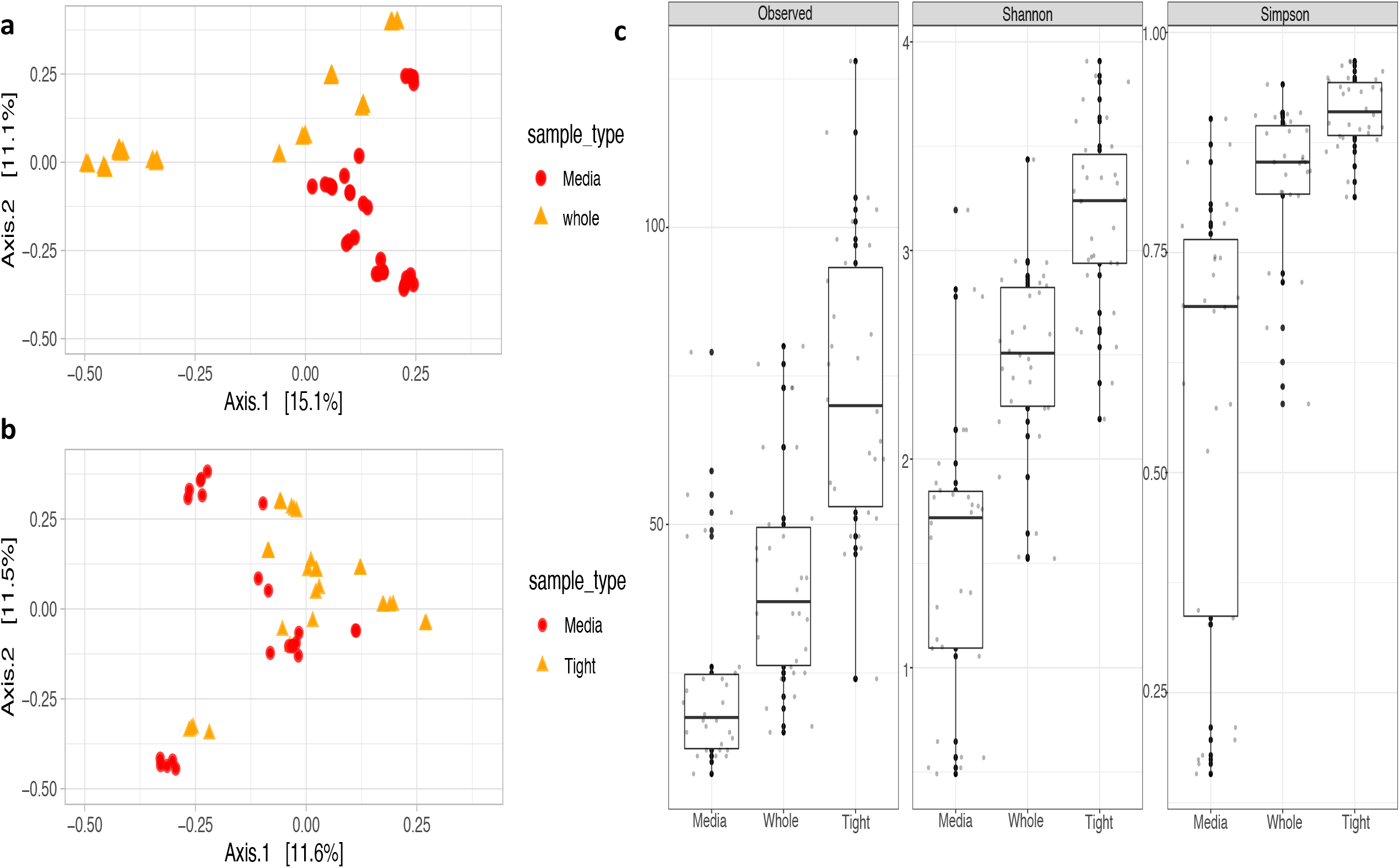
Principal coordinate analysis and alpha diversity to distinguish datasets. **a:** PCoA using Bray-Curtis dissimilarity to distinguish ‘media’ and ‘whole’ **b:** PCoA using Bray-Curtis dissimilarity to distinguish ‘media’ and ‘tight’. Colored dots/triangles represent replicate samples **c:** Alpha diversity differences between the three datasets: observed ASVs indicate richness, Shannon diversity index indicate overall alpha diversity and Simpson diversity index indicate evenness. Black dots represent the replicate samples. Median values are indicated by the horizontal line inside the box

In support of the beta diversity differences, clear differences in major taxonomic groups were observed between the media dataset and others. The media fraction of each strain consisted almost entirely of *Pseudomonadaceae* (Figure 2: a) accounting for more than 76% of the reads in each strain except VRM642 (45%). This was in stark contrast to the dominant groups for *Ostreobium*-associated datasets (*Methyloligellaceae* in ‘whole’ data and *Cyclobacteriaceae* in ‘tight’ data, Figure 2: b and c). These results imply that *Ostreobium*’s bacterial microbiome (bacteria that are tightly/loosely attached or intracellular) is distinct from loosely associated and unassociated microbiomes present in the media.

**Figure 2.**
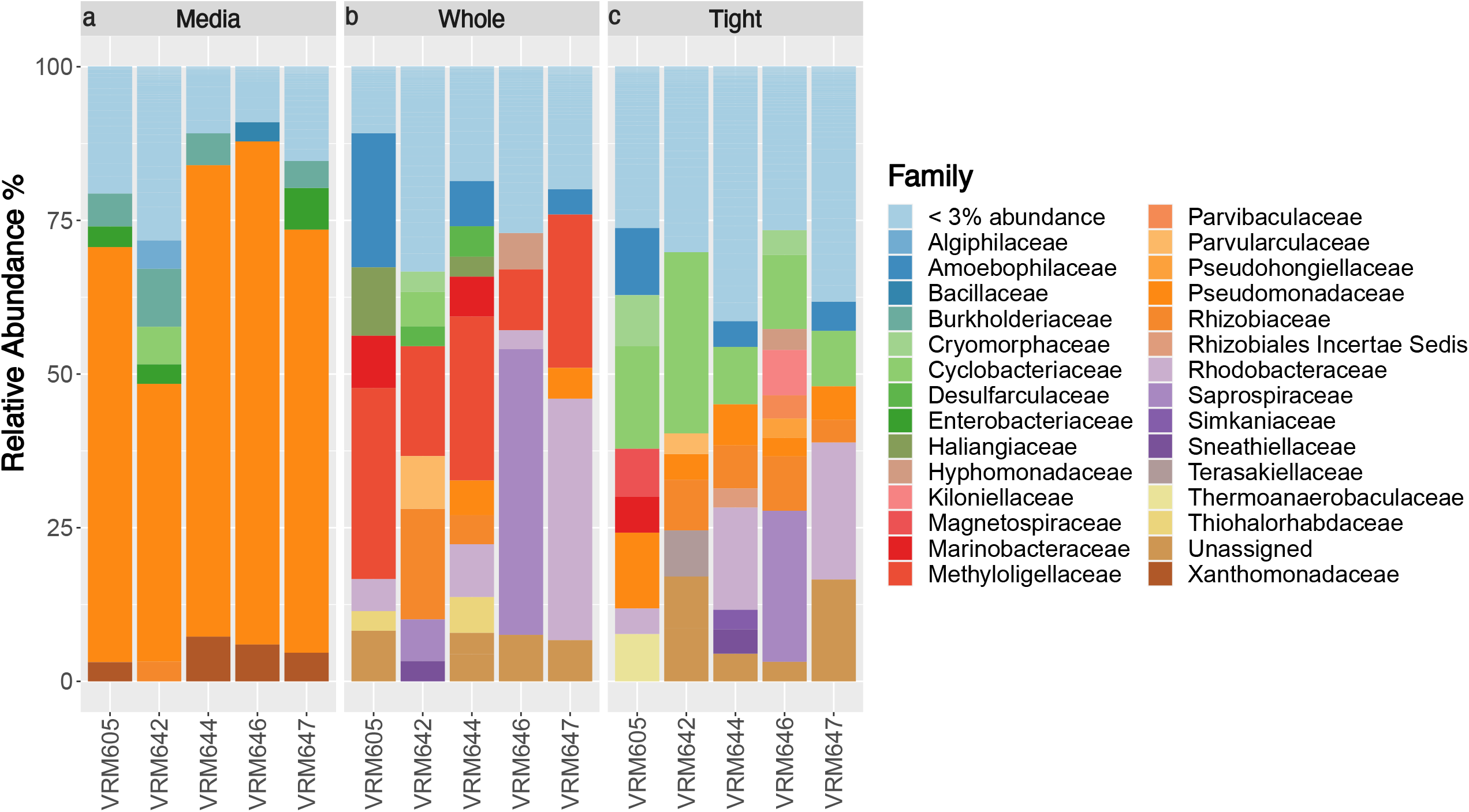
Bacterial community compositions of the three datasets: Relative abundances of major bacterial families found in ‘media’ (a), whole (b) and tight (c) datasets

### Taxonomically diverse bacterial associates and an unexpected methylotroph diversity

‘Whole’ data highlighted all the associated bacteria of *Ostreobium* while ‘tight’ data provided potential insights into those communities that are tightly attached and intracellular. In ‘whole’ data, *Proteobacteria* and *Bacteroidetes* were the most dominant phyla accounting for total relative abundance of 64.56% and 22.66%, respectively. The most dominant bacterial family (22.12%) was represented by a methylotrophic bacterium, *Methyloligellaceae* (c_*Alphaproteobacteria*, o_*Rhizobiales*, g_*Methyloceanibacter*) (Figure 2: b). About 37.85% of members were represented by *Rhodobacteraceae* (∼11.4%), *Saprospiraceae* (10.66%), *Amoebophilaceae* (7.01%), *Rhizobiaceae* (5.23%) and *Marinobacteraceae* (3.55%) (Supplementary data).

Of the 25 bacterial phyla detected in the ‘tight’ data, *Proteobacteria* (57.63%) and *Bacteroidetes* (31.43%) were the most dominant. Ninety different bacterial families were associated with *Ostreobium* strains and about 19.6% of the ASVs were taxonomically unidentified at family level. The dataset was dominated by the family *Cyclobacteriaceae* (15.3%) (Figure 2: c) which was only accounting for relative abundance of 2.36% in the ‘whole’ dataset. *Rhodobacteraceae* (9.56%), *Pseudomonadaceae* (6.34%) and *Rhizobiaceae* (5.7%) were the 2nd, 3rd and 4th most abundant families. *Cyclobacteriaceae* was represented by genera *Marinoscillum, Reichenbachiella, Ekhidna, Fabibacter* and *Fulvivirga*. At the genus level, *Leisingera* dominated the ‘tight’ dataset (Supplementary data file).

The most dominant family of the ‘whole’ dataset, *Methyloligellaceae*, was only contributing to 0.96% of the total relative abundance of the ‘tight’ dataset. A clear increase in *Pseudomonadaceae* in ‘tight’ data (6.34%) was observed compared to the ‘whole’ community (3.03%). The most abundant ASV of the ‘tight’ data was represented by an uncultured *Alphaproteobacteria* followed by ASVs representing *Pseudomonadaceae* (g_Pseudomonas) and *Rhodobacteraceae* (g_ *Leisingera*). *Chlamydiae*, typically an intracellular taxon, was also a highly abundant phylum (∼1.35%) in the’ ‘tight’ data while only accounting for 0.27% of the reads in the ‘whole’ dataset. There was a clear increase in relative abundance of *Kiloniellaceae* (2.28%) in ‘tight’ compared to ‘whole’ data (0.74%). In both datasets, *Leisingera* was found to be highly abundant, ranking as the most abundant in the ‘tight’ dataset as well as the second most abundant in the ‘whole’ dataset (Supplementary data file).

Since the ‘whole’ dataset was dominated by a methylotrophic bacterium, both ‘whole’ and ‘tight’ datasets were inspected for the presence of other methylotrophs, yielding several other methylotrophic genera such as *OM43* (f_ *Methylophilacea*), *Methylophaga* (f_ *Methylophagaceae*), *Leisingera* (f_ *Rhodobacteraceae*) and *Filomicrobium* (f_*Hyphomicrobiaceae*) [58]–[60] in varying relative abundances. Among them, *Leisingera* was consistently associated with all strains in both ‘whole’ and ‘tight’ datasets.

### Closely associated bacteria of *Ostreobium*

The protocol employed in the generation of the ‘tight’ data provided an opportunity to study an enriched fraction of attached and intracellular bacteria which may be closely associated and possibly influence *Ostreobium* physiology. Alpha diversity as assessed by observed ASV richness, and Simpson and Shannon indices showed that the number of bacterial taxa associated and their evenness were similar among the strains, with none of the differences statistically significantly (Kruskal-Wallis test: chi-squared _Observed (1)_ = 8.0383, p =0.09018, chi-squared _Simpson (1)_ =8.0473, p =0.08986, chi-squared _Shannon (1)_ =6.8516, p =0.1439).

The bacterial community compositions were distinct among the *Ostreobium* strains, with a statistically significant effect of strains on bacterial community composition inferred for Bray-Curtis distances (p=0.001, F.Model=4.4042, R^2^=0.41338), phylogenetically aware distances, Unifrac (p=0.001, F.model=5.2818, R^2^=0.45802) and weighted Unifrac (p=0.001, F.model=7.02, R^2^=0.52901). The analysis of multivariate homogeneity of group dispersions were insignificant for Bray Curtis and Unifrac distances confirming that *Ostreobium* strains have significantly different communities. Considering these strains represent five different *Ostreobium* clades, these results indicate that the *Ostreobium* microbiome may be impacted by phylogeny.

*Ostreobium* strains were consistently associated with certain bacterial groups indicating the presence of a core microbiome, with core phyla including *Acidobacteria, Actinobacteria, Bacteroidetes, Gemmatimonadetes, PAUC34f, Planctomycetes, Proteobacteria* and *Verrucomicrobia*. The core families were represented by 34 bacterial families (including some unassigned at the family but consistently associated, for example: *Actinomarinales* unassigned at family level) (Figure: 3) accounting for over 60% of reads in all strains and even >75% in four out of five strains. However, at the ASV level, only 6 ASVs (represented by Pseudomonadaceae (2 ASVs), *Marinobacteraceae, Sneathiellaceae, Rhizobiaceae* and an *Actinomarinales* unassigned at family level) were consistently associated among all the strains averaging for ∼10% of the total reads. This implies that although *Ostreobium* associates with conserved bacterial families their members are different at a finer taxonomic level possibly representing different species or strains. This was also clear from the presence/absence of the ASVs representing the 34 core families across the strains (Table S1). These results imply that the community structure difference across the strains is mainly driven by different members of the core bacterial families being hosted by different *Ostreobium* strains.

**Figure 3.**
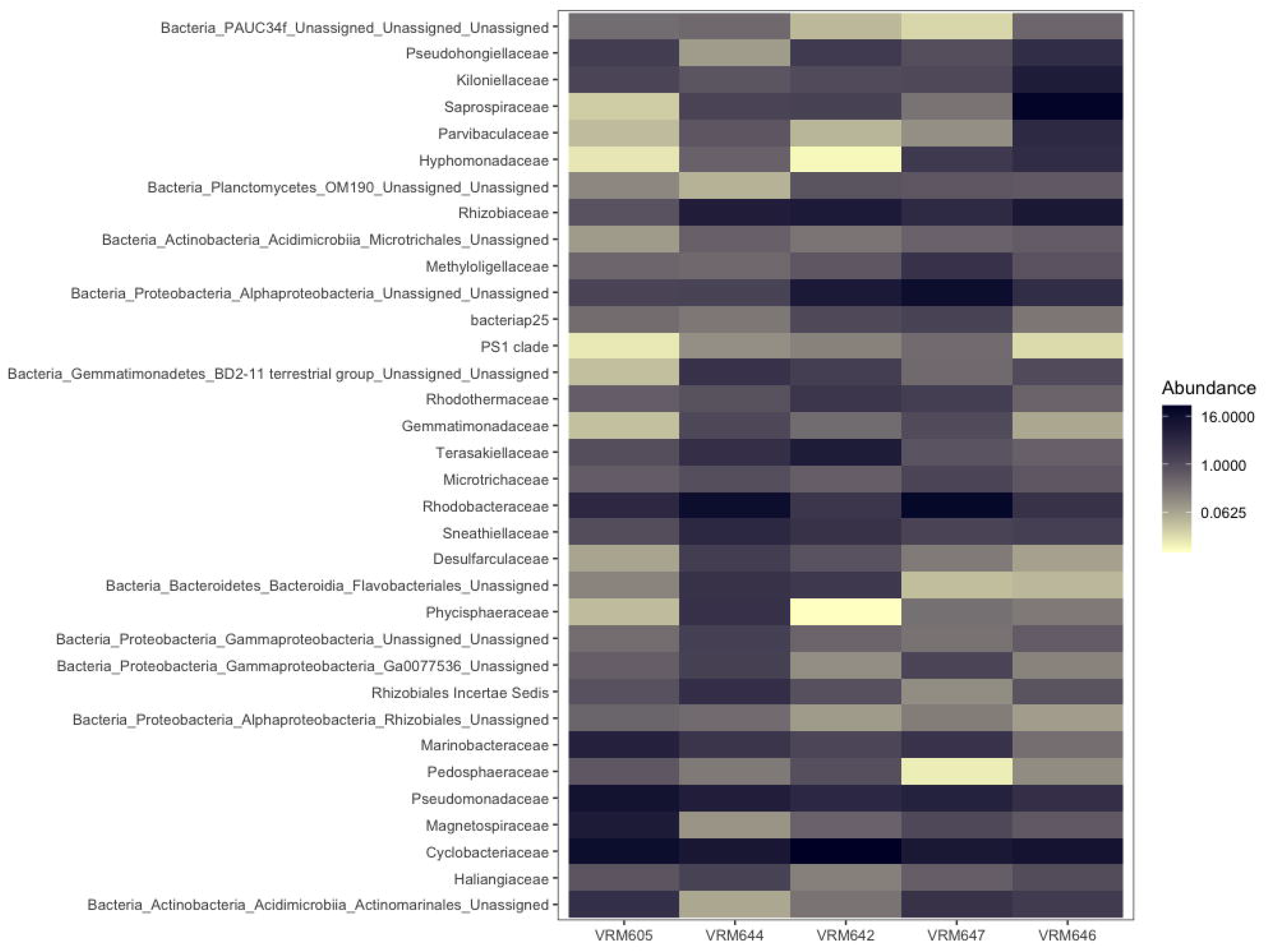
Heat map of the core bacterial families: The relative abundance of the 34 core bacterial families detected using the ‘tight’ data across different strains

Next, we attempted to determine the likely physical associations through an indicator taxa analysis (Table S2). Indicators of ‘tight’ data could be thought of as those that are potentially tightly attached or intracellular. As it is expected for the loosely attached communities to decrease in abundance following the serial washing step. Likewise, indicators of ‘whole’ data could be thought of as those that are loosely attached. Among the 39 bacterial taxa detected as indicators of ‘tight’ fraction, *Cyclobacteriacea* (stat = 0.594, p =0.001) and members of candidate phylum PAUC34f (stat =0.614, p=0.001) were identified as the most strongly associated. Both these groups were previously identified as core taxa of *Ostreobium*. Other than these two, 15 more core bacterial families such as *Pseudohongiellaceae, Alphaprotoeobacteria* unassigned at family level, *Rhodothermaceae, Pedosphaeraceae, Kiloniellaceae, Gemmatimonadaceae, bacteriap25, Rhizobiales Incertae Sedis, Terasakiellaceae, Magnetospiraceae, Parvibaculaceae, Phycisphaeracea, Gammaproteobacteria* unassigned at family level and *Flavobacteriales* unassigned at family level were also found as indicators of the ‘tight’ data. *Methyloligellaceae*, another core taxa (stat = 0.878, p =0.001) was the most strongly associated family of the ‘whole’ data out of the 13 indicator taxa. Indicators of the ‘whole’ data also represented 5 more core bacterial families such as *Actinomarinales* unassigned at family level, *Microtrichaceae, Mitrotichales* unassigned at family level, *Haliangiaceae* and *Desulfarculaceae*. The indicator taxa analysis also found 14 taxa (table S2) associated with both the ‘tight’ and ‘whole’ dataset. These taxa also represented core bacteria taxa such as *Saprospiraceae, Gemmatimonadetes* unclassified a family level, *OM190* unclassified at family level, *Hyphomonadaceae, Rhizobiaceae, Rhodobacteraceae, Sneathiellaceae* and *Marinobacteraceae*. These taxa could also be thought as those that are potentially tightly attached or intracellular. In total, 32 core bacterial taxa were found as indicators of either ‘tight’, ‘whole’ or ‘tight and whole’. These results indicate that the majority of the core taxa of *Ostreobium* may have an attached lifestyle. *Pseudomonadaceae* (stat =0.889, p = 0.001) was the most strongly associated family of the ‘media’ data.

### Culture-based associations are observed in the co-occurrence networks of *Ostreobium* and bacteria

The co-occurrence network created using *P. lutea* skeletal samples consisted of 138 nodes and 322 edges while the *P. australensis* co-occurrence network consisted of 85 nodes and 209 edges. The global network properties for each network can be found in Supplementary table S3. By clustering, we identified one *Ostreobium*-bacterial module from *P. lutea* and one from *P. australensis* (Figure 4: a and b). The *P. australensis* module consisted of 2 representative *Ostreobium* ASVs annotated as *Ostreobiaceae*. This module included 35 nodes and 104 edges (Figure 4: a). The *P. lutea* module consisted of 1 representative *Ostreobium* ASV (annotated as *Ostreobiaceae*) and included 126 nodes and 310 edges (Figure 4: b). Both *Ostreobium*-bacteria modules showed high clustering coefficients (> 0.5) indicating densely connected neighbourhoods.

**Figure 4.**
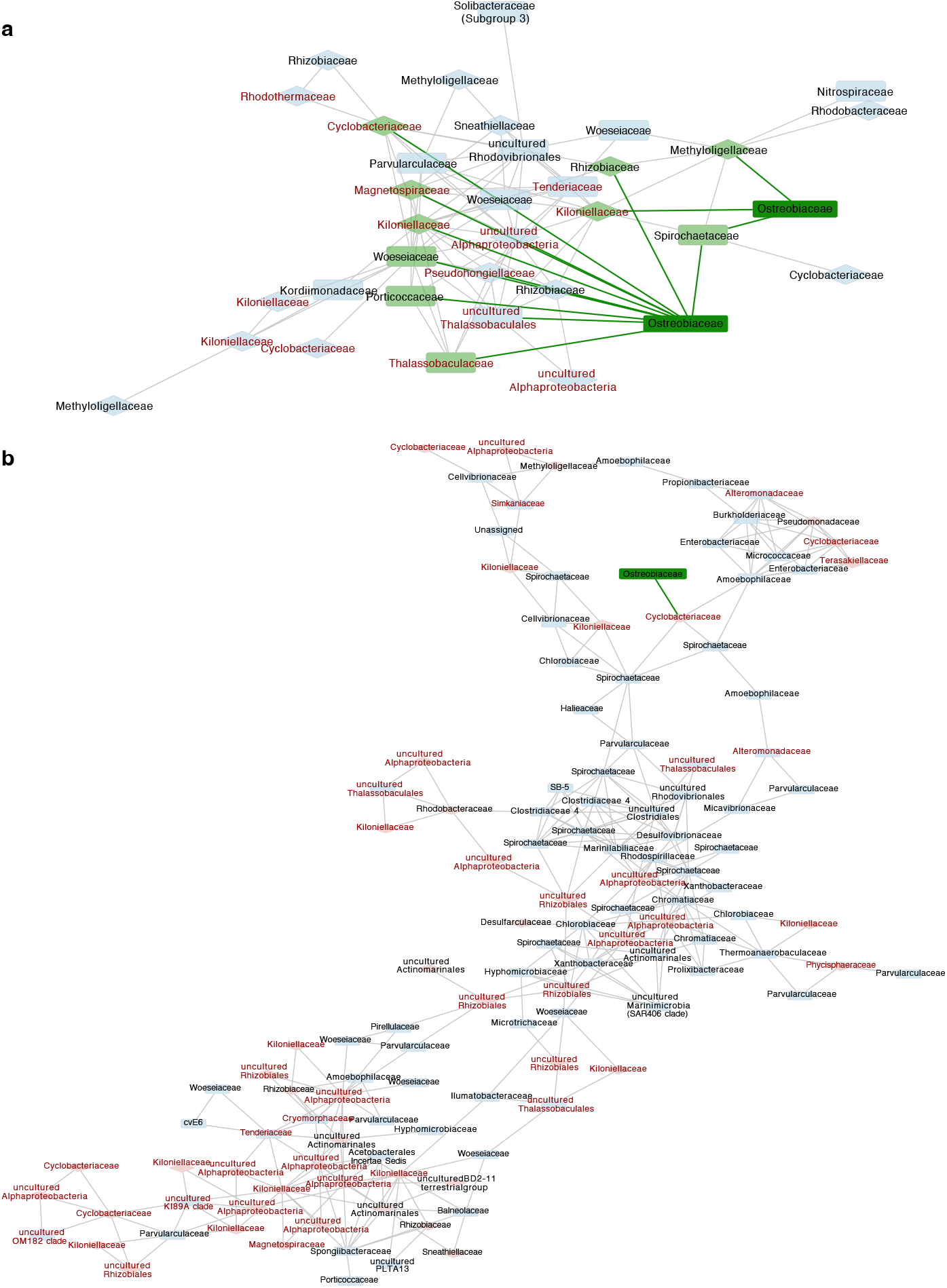
Co-occurrence network analysis of *Paragoniastrea australensis* and *Porites lutea* skeletal samples: *Ostreobium*-bacteria module from *P. australensis* (a) and *P. lutea* (b). Diamond shape nodes indicate core bacterial families of ‘tight’ data. Taxa labelled in red represent those that were found as indicators of ‘tight’ data. Direct edges from *Ostreobium* nodes are colored in green. Edge width is continuously mapped to edge weight

The most important observation was that the majority of bacterial communities potentially interacting with *Ostreobium* in the natural environment represented most of the core bacterial families (Figure 4: diamond shaped nodes) and indicator taxa of ‘tight’ data (Figure 4: taxa labelled in red). For instance, in *P. australensis*, representative *Ostreobium* nodes showed significant co-occurrences with many *Cyclobacteriaceae* and *Kiloniellaceae* (Figure 4: a) which were found to be both core and indicator taxa of the ‘tight’ data. In *P. lutea*, the representative *Ostreobium* node was connected to the rest of the community via *Cyclobacteriaceae* (Figure 4: b) highlighting possible tight association with this bacterial lineage.

### *Ostreobium* phylogeny correlates with bacterial community structure

A significant difference in community structures among strains led us to investigate patterns of phylosymbiosis. We expected the *Ostreobium* phylogeny to have significant congruence with the microbial dendrograms. We first quantified the phylosymbiotic signal using the ‘whole’ dataset representing all the culture associated bacteria and found no statistically significant congruence between the trees (nMC = 0.6, p=0.8755 for all Bray-Curtis, Unifrac and weighted Unifrac UPGMA comparisons with the host tree). For the ‘tight’ data, on the other hand, topological congruence analysis showed a significant (p=0.0272) association between the host phylogenetic tree and microbial dendrogram constructed on Bray Curtis distances, with a normalised matching Cluster score of 0.3 (Figure: 5 and Table 1). Normalized Matching Cluster scores closer to zero indicate higher topological congruence and reveal signatures of phylosymbiosis. With over 60% of reads in each strain stemming from ASVs representing core bacterial families (see above), we investigated whether they were responsible for the significant signal observed and we detected the same significant nMC of 0.3 for the core microbial Bray Curtis distances. Although core weighted and unweighted UniFrac distances generated non-significant results, more evidence for phylosymbiosis was apparent based on improved p values compared to the previous analysis on the entire community (Table 1).

**Figure 5.**
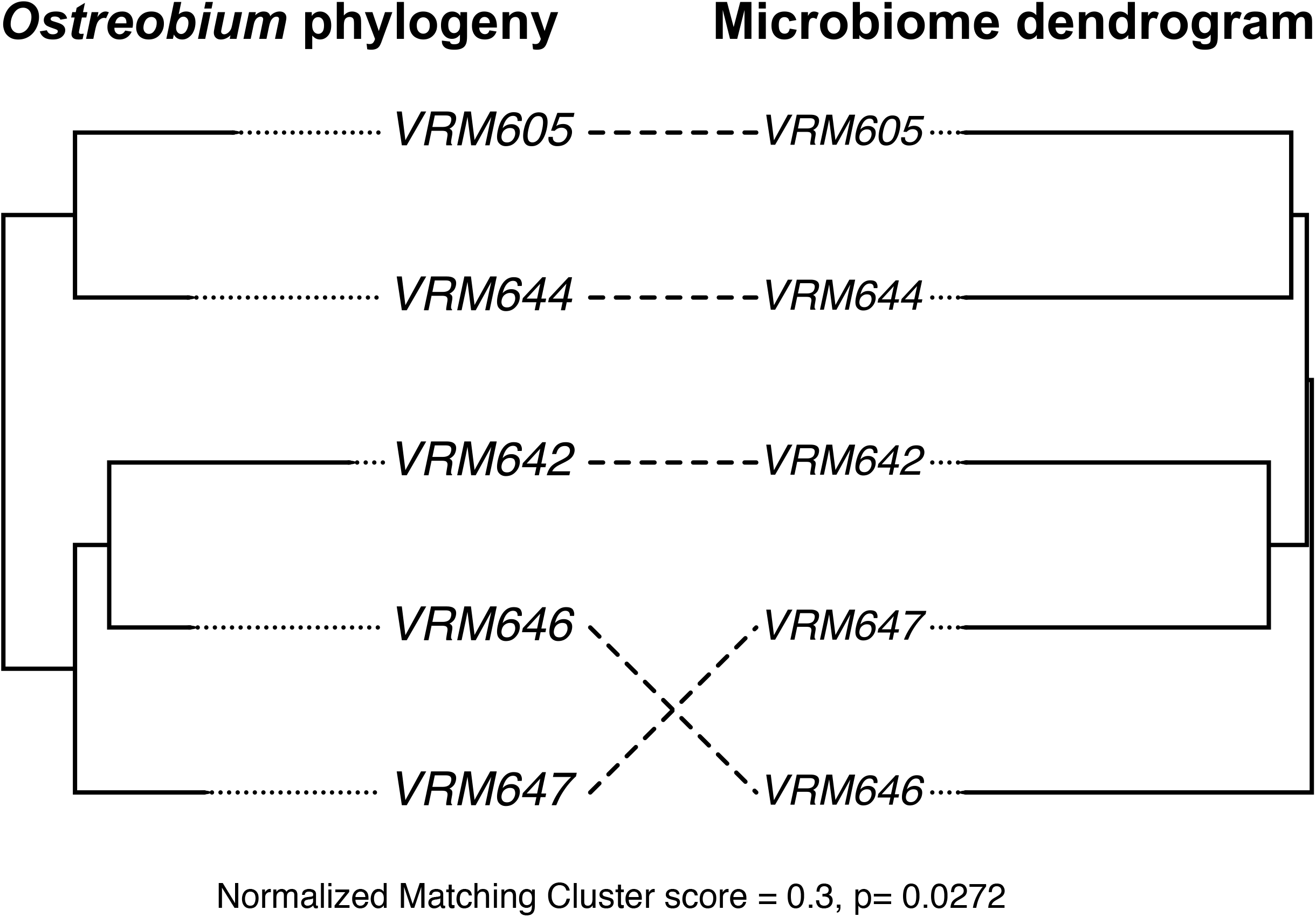
Topological congruence between the host phylogeny and microbial dendrogram: Host phylogenetic tree was constructed using *tufA* sequences. The microbiome tree represents the UPGMA tree constructed on Bray-Curtis distances. This was built using the ASVs representing the core bacterial families of the ‘tight’ data. The normalised matching cluster score (nMC) and p value indicates significant congruence

**Table 1:** Summary of Phylosymbiosis testing: ‘Testing type’ highlights different tests carried out using the ‘tight’ data. Normalised matching score ranges from 0 (complete congruence) to 1 (complete incongruence).

| Testing type | Distance metric | Normalized matching cluster score | p value |
| --- | --- | --- | --- |
| Entire community | **Bray** | **0.3** | **0.0272** |
| Entire community | Unifrac | 0.6 | 0.8687 |
| Entire community | Weighted-Unifrac | 0.6 | 0.8764 |
| Core (ASVs representing core bacterial families) | **Bray** | **0.3** | **0.0272** |
| Core (ASVs representing core bacterial families) | Unifrac | 0.4 | 0.1657 |
| Core (ASVs representing core bacterial families) | Weighted-Unifrac | 0.4 | 0.1687 |

## Discussion

Despite extensive work on the coral holobiont, many questions about the major algal symbiont of coral skeleton, *Ostreobium*, remain to be answered. Understanding how the major algal symbiont of the coral interacts with the surrounding microbiome helps to extend our knowledge on the coral holobiont. As a first step towards understanding these interactions, we describe the bacterial microbiome of *Ostreobium* using diverse clades of cultured *Ostreobium*. This study led to the identification of taxa that were likely to be intracellular or closely attached, which can be used to guide future studies to confirm their location. Our study revealed that *Ostreobium* consistently associates with 34 bacterial taxa, irrespective of strain, that constitute the majority of its microbiome. However, at a finer taxonomic resolution, these bacterial families are mostly represented by distinct ASVs in each strain leading to significant community differences between the strains. We identified phylosymbiotic signatures stemming from these core bacterial families implying that they may preferentially associate or co-differentiate with *Ostreobium* hosts. By constructing co-occurrence networks on coral skeletal samples, we show that the culture-based associations inferred in this study exist in natural settings and the taxa we identified as potentially closely associated frequently co-occur with *Ostreobium*.

The *Ostreobium* microbiome included bacteria that have been previously found in coral skeleton microbiome. Bacteria representing *Candidatus Amoebophilus, Kiloniellaceae, Rhizobiales* and *Myxococcales* were recently shown to be preferentially associated with the coral skeleton microbiome [61]. We detected these bacterial taxa in high abundance in *Ostreobium* cultures, suggesting that their presence in the coral skeleton may be due to an association with *Ostreobium*. The *Cyclobacteriacea*, previously shown as an unusual suspect consistently associated with corals [61],was here shown to be a potentially tightly attached and perhaps even intracellular bacterium of *Ostreobium*. In addition to these, the microbiome found in our study was dominated by bacterial phyla predominantly found in coral skeletons, such as *Proteobacteria, Bacteroidetes, Actinobacteria, Planctomycetes* and *Acidobacteria* [62]–[64]. We also found a significant abundance of *Chlamydia* which was shown to be potentially involved in symbiosis with the eukaryotes in the skeleton through a metagenomic analysis [65]. Moreover, consistent with observations from Ricci et al [35] on coral skeletal samples, we also found an ASV assigned to an uncultured bacterium from phylum *Actinobacteria* representing class *Acidimicrobiia* in all the *Ostreobium* strains.

Our three-dataset study design allowed distinguishing the host associated microbiome from the taxa in the media fractions and investigating their likely physical associations. The media fractions of each strain were less diverse, almost entirely composed of *Pseudomonadaceae* and were significantly different in composition from the host associated data. Although our data do not constitute proof of physical associations, they shed light on likely locations of particular bacterial groups. In generating our ‘tight’ data, our approach was to wash away most of the loosely attached and contaminating bacteria through serial washing, enriching tightly associated bacteria. We cut open the filaments and collected the cytoplasmic material to further enrich intracellular bacteria. Unexpectedly, the ‘tight’ data showed the highest richness. This may be attributed to the methodology we used for data generation. It is likely that cutting open the filaments has revealed bacteria that were not detected in the ‘whole’ dataset due to being present at low abundance compared to the loosely attached bacteria and those found in the media. While none of these methods confirm the nature of physical association, they do show trends reflecting which bacteria are more likely to be tightly associated with their host.

Comparison between ‘tight’ with ‘whole’ data allowed identifying taxa that are more likely to be intracellular or tightly attached. The *Methyloligellaceae*, a very abundant bacterial family in the ‘whole’ dataset, was significantly reduced in the ‘tight’ dataset, suggesting that the serial washing may have largely removed these taxa, which we could therefore speculate to be loosely attached to the host algal cell wall. Both the taxonomic composition analysis and indicator taxa showed *Cyclobacteriacea* to be significantly associated with the ‘tight’ data implying they may be closely attached or potentially intracellular. *Cyclobacteriaceae* are known to degrade polysaccharides which could contribute to the carbon metabolism of the coral holobiont. Moreover, they are involved in carotenoid biosynthesis, antibiotic resistance, and quorum-sensing, all of which can benefit the alga [66]. *PAU34f*, a candidate phylum known for its symbiotic associations of sponges, was also found to be significantly more prevalent in the ‘tight’ dataset. Genomic studies have revealed that they have the potential to degrade both sponge and algae derived carbohydrates, presumably provide phosphate reservoirs to the sponge host in deprivation periods, produce antimicrobial compounds that can be used by the host as a defence strategy and harbour signatures of host associations in the genome [67].

Our results show that *Ostreobium* is colonised by a core set of bacterial families that differ on a finer taxonomic scale, resulting in differences in the community structure among strains. By studying these core bacterial taxa, we identified key functional types associated with *Ostreobium*. Most of the bacteria represented known nitrogen cyclers (*Rhizobiaceae, Terasakiellaceae, Kiloniellaceae, Sneathiellaceae* [68]–[71], Sulphate reducers (*Desulfarculaceae* [72], polysaccharide degraders (*Cyclobacteriaceae, PAUC34f, Pseudohongiellaceae* [66], [67], [73] and antimicrobial compound producers (*Cyclobacteriaceae, Myxococcales* (families *bacteriap25* and *Haliangiaceae* [66], [74]–[76]). Apart from these, we also found potential vitamin B12 producers such *Rhodobacteraceae* (*Roseobacter* clade) which are known mutualists of eukaryotes [1], [77]. *Ostreobium*’s dependence on bacteria for vitamin B12 was previously shown and it was hypothesised based on metatranscriptomic data of the coral holobiont that *Rhodobacteraceae* may provide this vitamin [78]. Members of *Rhizobiaceae* are nitrogen fixers, and this metabolism has also been documented in *Terasakiellaceae* [68]. Both *Kiloniellaceae* and *Sneathiellaceae* are potential denitrifiers involved in reduction of nitrates to gaseous nitrogen [69], [71]. *Kiloniellaceae* were previously shown to be closely associated with *Symbiodinaceae* [79] and a preferential coloniser of the coral skeleton [61]. Our results and previous studies therefore suggest that *Kiloniellaceae* may be an important associate of both algal symbionts of the tissue and skeleton. Nitrogen fixation can benefit the alga by providing an organic nitrogen source and also the coral holobiont, which lives in an oligotrophic environment by contributing to the nitrogen budget [80]. The presence of both nitrogen fixers and denitrifiers shows that the *Ostreobium*-associated microbiome may contribute to nitrogen homeostasis, contributing to the stabilisation of the coral holobiont. Moreover, our indicator analysis revealed a range of bacterial taxa as indicators of ‘tight’ data, in addition to *Cyclobacteriaceae* and *PAU34f*. In total, out of the 34 core bacterial families, 32 of them were either indicators of ‘tight’, ‘whole’ or ‘tight and whole’. These results show that the majority of the core bacterial families of *Ostreobium* may live attached (tightly or loosely) to or potentially inside the *Ostreobium* filaments. Additional work should be carried out to confirm these associations through 3-dimensional imaging.

The associations of *Ostreobium* with a range of methylotrophic bacteria is intriguing. Methylotrophs can utilise reduced carbon substrates without carbon-carbon bonds (i.e. C1 substrates) as their source of carbon and energy [59]. Methanotrophs, the methylotrophs that can utilise the potent greenhouse gas methanol are of special interest with regard to climate change [81]. While we detected diverse methylotrophs such as *OM43* (family *Methylophilaceae*), *Methylophaga* (family *Methylophagaceae*) and *Filomicrobium* (family *Hyphomicrobiaceae*) in the microbiome, the genus *Methyloceanibacter* of family *Methyloligellaceae* and *Leisingera* of family *Rhodobacteraceae* were the most dominant and consistently associated. Members of the family *Methyloligellaceae* have been shown to utilise both methylated compounds and methane as their carbon source [82]. *Leisingera*, represent organisms that can grow by oxidation of methyl groups and specifically *L*.*methylohalidivorans* use methyl halides as the sole source of carbon and energy [83], [84]. Interestingly, *Leisingera* species have also been shown to produce antimicrobial compounds and secondary metabolites such as siderophores and acyl-homoserine lactones involved with quorum sensing that may aid in symbiotic relationships [85]. The fact that methylotrophs are commonly isolated from macroalgae [86] suggests unsuspected algae-methylotroph associations. Considering the diversity and consistent association of some methylotrophs with *Ostreobium*, our results suggest a possible production of methanol and/or methylated compounds by *Ostreobium*. Recently, methylotrophic genus *Methylobacterium* was identified as an intracellular core genus of *Symbiodinaceae* [79]. These findings suggest possible intricate relationships between methylotrophs and algal symbionts of the coral organism. A relationship that may be similar to terrestrial plants and methylotrophs where these bacteria affect the overall health of the holobiont by producing plant growth hormones [87]. Overall, *Ostreobium* associated methylotrophs may play an important role in C1 metabolism of the holobiont.

Our co-occurrence network analysis revealed two *Ostreobium*-bacteria modules from both coral species, *P. lutea* and *P. australensis*. Network modules are thought to represent biologically meaningful microbial communities [88]. Bacterial taxa in both these modules represented most of the core bacterial families and indicators of ‘tight’ data. As described earlier, we proposed these core and indicator taxa as the closely associated microbiome of *Ostreobium* that may be potentially influencing the *Ostreobium* physiology. Co-occurrence results suggest that *Ostreobium* tends to maintain these important associations even in cultures. These significant co-occurrences help to further support our hypothesis. Overall, we demonstrated that cultured *Ostreobium* harbours bacteria that have been previously described in coral skeleton microbiomes and those that co-occur significantly in the natural coral environment. These results show that culture-associated microbiota represents associations present in their natural environment.

Since the beta diversity analysis on ‘tight’ data showed a significant effect of host lineage on community composition, it led us to quantify the correlation between the host phylogeny and bacterial community composition to look for patterns of phylosymbiosis. We quantified the phylosymbiotic signal from both ‘whole’ and ‘tight’ data to investigate how the signal differs between the two. Our results indicated that the signal from all the culture-associated bacteria in ‘whole’ were insignificant. While phylosymbioses involve trends across the entire microbiome composition, their absence does not rule out that bacteria preferentially associate with certain species of hosts or co-differentiate with them [89]. By analysing the closely associated microbiome using the ‘tight’ dataset, we found that the microbial dendrograms built on the entire community composition as well as the ASVs representing the core bacterial families produce the same significant results highlighting the origination of the phylosymbiotic signal. Consequently, we hypothesise that *Ostreobium* preferentially associates with these core bacterial taxa and some of these bacteria may be evolutionarily conserved.

Overall, we provide a comprehensive study on the microbiome of the major coral skeleton symbiont *Ostreobium* and shed light on phylosymbiotic signatures of an algal-bacterial system. Our results indicate preferential associations between certain bacterial taxa and *Ostreobium* which warrants further testing of these associations both from functional and evolutionary perspectives. Our findings extend the knowledge on the microbiome of endolithic algae, coral holobiont, coral reef microbial ecology and enhances our understanding of evolutionary relationships between microalgae and bacteria.

## Acknowledgments

We are grateful for Stephen Wilcox from Walter and Eliza Hall Institute of Medical Research for his assistance in library sequencing. We thank Dr. Francessco Ricci for providing raw QIIME2 files linked to Bioproject PRJNA753944 and Dr. Cintia Iha for providing valuable help in Bioinformatic analysis.

## Author contributions

UP, HV, AW and KT designed the research. UP performed DNA extraction, library preparation and bioinformatic analyses. UP wrote the first draft of the manuscript. All authors contributed to the final edited version of the manuscript.

## Data availability

Sequence data are available under NCBI BioProject ID PRJNA910678

## Supplementary

**Table S1:** Amplicon Sequence Variants representing the 34 core bacterial families detected in ‘tight’ data and their prevalence across the strains

**Table S2:** Indicator species analysis between the three datasets to identify bacterial families that preferentially associate a particular dataset

**Table S3:** Global network properties of *Paragoniastrea australensis* and *Porites lutea* co-occurrence networks

## Supplementary data file

